# Accelerated detection of Gram negative bacteria in blood culture by enhanced acoustic flow cytometry (AFC) following peptide nucleic acid fluorescence *in situ* hybridization (PNA-FISH)

**DOI:** 10.1101/370171

**Authors:** Xiao Xuan Huang, Nadezda Urosevic, Timothy J.J. Inglis

## Abstract

Bacteraemia is a risk factor for subsequent clinical deterioration and death. Current reliance on culture-based methods for detection of bacteraemia delays identification and assessment of this risk until after the optimal period for positively impacting treatment decisions has passed. Therefore, a method for rapid detection and identification of bacterial infection in the peripheral bloodstream in acutely ill patients is crucial for improved patient survival through earlier targeted antibiotic treatment. The turnaround time for current clinical laboratory methods ranges from 12 to 48 hours, emphasizing the need for a faster diagnostic test. Here we describe a novel assay for accelerated detection of bacterial infection in blood culture (BC) using peptide nucleic acid fluorescence *in situ* hybridization enhanced acoustic flow cytometry (PNA-FISH-AFC). For assay development, we used simulated blood cultures (BCs) spiked with one of three bacterial species: *Escherichia coli, Klebsiella pneumoniae* or *Pseudomonas aeruginosa* at a low concentration of 10 CFU/mL. Under current clinical settings, it takes a minimum of 12 hours incubation to reach positivity on the BacTEC system, corresponding to a bacterial concentration of 10^7^-10^9^ CFU/mL optimal for further analyses. In contrast, our PNA-FISH-AFC assay detected 10^3^ – 10^4^ CFU/mL bacteria in BC following a much shorter incubation of 5 to 10 hours in culture. Using either PCR-based FilmArray^®^ assay or MALDI-TOF for bacterial detection, it took 7-10 and 12-24 hours of incubation, respectively, to reach the positive result. These findings indicate a potential time advantage of PNA-FISH-AFC assay over currently used laboratory techniques for rapid bacterial detection in BC with significantly improved turnaround time.

## Introduction

Sepsis is one of the leading causes of death from infection, usually accompanied by bacteraemia. In its most fulminant form sepsis has a mortality rate of up to 40-60% [1-3]. Sepsis-associated bacteraemia may be mono- or polymicrobial and requires lengthy laboratory processes to identify and characterize the causative agent(s). The time required for initial bacterial detection in the diagnostic laboratory ranges from 12 to 48 hours and depends in part on bacterial growth rate and the number of culture-dependent laboratory procedures required after initial isolation.

The standard method for detection of bacteremia boosts bacterial cell numbers by culture of peripheral blood (blood culture, BC) to aid detection and subsequent analysis. To do this, an 8-10mL sample of patient’s blood is aseptically inoculated into each of a pair of aerobic and anaerobic BC bottles that contain bacterial growth media, lytic agents to release phagocytosed bacteria from leukocytes and ion exchange resin beads to neutralize the effect of residual antibiotics and improve bacterial recovery [4]. Once inoculated BCs reach the clinical laboratory, they are incubated in a dedicated autoanalyser that continuously monitors changes in pH or carbon dioxide production as indicators of bacterial growth [5]. The autoanalyser signals the BC bottles need further analysis after reaching a pre-set detection threshold. The positive signal threshold is set to correspond to the minimum point at which Gram stain can be used to reliably confirm the presence of bacteria and direct downstream analyses.

Growth rates of bacteria vary greatly in BCs. Fast growing bacteria such as *Escherichia coli* and *Klebsiella pneumoniae* may reach the detection threshold in shorter periods of time than slower growers such as *Pseudomonas aeruginosa*, *Staphylococcus aureus* and *Streptococcus pyogenes* that may require 19 to 48 hours, or fastidious organisms that can take as long as up to five days. The initial bacterial load in blood plays a critical role in determining time to detection in BC. The bacterial load in the blood of bacteraemic adult patients is typically very low, ranging between 1 to 10 CFU/mL, but in more severe cases may exceed 10^3^ CFU/mL. There is a strong correlation between bacterial loads in patients’ blood, time to positivity in BC, increased in-hospital stays and mortality in bacteraemic patients [6-9]. This has prompted further development of rapid diagnostic tests to reduce the time taken to detect, identify and perform antimicrobial susceptibility testing in order to optimize antibiotic treatment and improve patients’ survival.

On detection of a positive BC, aliquots are usually taken for initial presumptive and subsequent definitive identification tests. Three main options for bacterial identification are in current use: (1) testing directly from blood culture bottles; (2) sub-culture to obtain single colony growth; (3) testing after enrichment and subsequent purification (Fig 1). Each of these may use nucleic acid-based identification processes (e.g. probe-based fluorescence microscopy, microarray, PCR, DNA sequencing), protein characterization by matrix-assisted laser desorption/ionization time-of-flight (MALDI-TOF) mass spectrometry, or substrate utilization methods (e.g. VITEK or API), respectively. The final bacterial identification may take an additional 2 to 36 hours for a definitive result, depending on the bacterial species and range of tests performed. Currently, the most rapid turnaround time from BC inoculation to definitive bacterial identification is approximately 16 hours using molecular techniques, while conventional sub-culture based identification may take an additional 48 hours or more [10].

**Fig 1.**
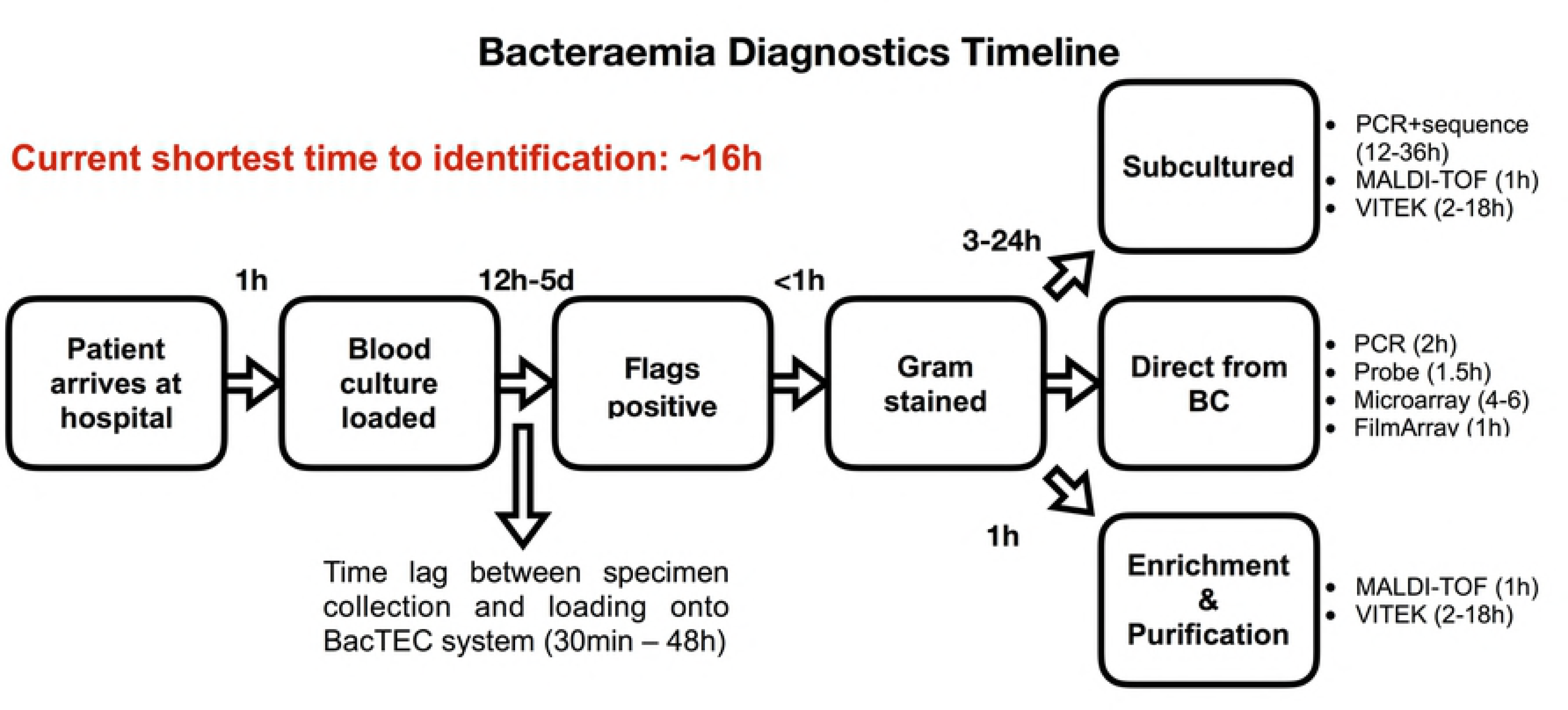
Timeline of current bacteraemia diagnostics. **D**iagram showing major stages in current bacteraemia diagnostics featuring three main test streams with a minimum of 16 hours and maximum of 2 or more days for definitive bacterial identification.

The two rapid, culture-dependent technologies currently used in the laboratory confirmation of bacteraemia are MALDI-TOF mass spectrometry and PCR assays. Although robust enough for use in clinical laboratories, these technologies have shortcomings that may result in failure to detect the target or otherwise misleading data. The minimal bacterial concentration for microbial identification by MALDI-TOF is above 10^7^-10^8^ CFU/mL for reliable genus or species identifying scores in the range of 1.7-2.0 [11-13]. Bacterial concentrations lower than 10^7^ CFU/mL are more likely to generate unreliable identification due to a low signal-to-noise ratio caused by background of media and blood elements. Careful MALDI-TOF result interpretation is required with polymicrobial blood cultures where the error rate can be three times higher [10] and only predominant microorganisms could be identified [12]. Gram negative bacteria are easier to detect in polymicrobial BCs than Gram positive organisms due to their rapid growth rate [14]. MALDI-TOF performs poorly on Gram positive rods and yeasts due to their slow growth, resulting in a low microbial concentration [10]. Alternatively, while molecular detection by PCR can give specific results with a lower concentration of bacteria than MALDI-TOF, the sensitivity of detection is adversely affected by the sample matrix. In complex systems containing blood, urine and sputum, components such as haemoglobin can inhibit the PCR and cause false negative results [15]. In addition, the presence of microbial genetic material as demonstrated by PCR assay does not guarantee the presence of underlying bloodstream infection since it can originate from distant focal points in other tissues. This and carry-over contamination from laboratory equipment or reagents are common causes of false positive results, since a 16S rRNA gene-based detection probe can detect eubacterial DNA found ubiquitously in the environment [16]. Consequently, there is a need to confirm true bacteraemia by specific detection of intact bacterial cells.

Peptide nucleic acid (PNA) fluorescence *in situ* hybridization (FISH) has been used extensively in fluorescence microscopy to study pure growth on solid media, identify bacteria and yeast in liquid culture media and in positive blood cultures [17-19]. The repeated units of N-(2-aminoethyl)-glycine give the PNA probe a neutral backbone [20-21] making it more stable and resistant to enzymatic and pH degradation. This, in turn, allows strong binding to DNA and RNA targets under low ionic conditions and easier access to secondary RNA structures, making PNA probes specific, more stable and a better alternative to oligonucleotide probes.

Flow cytometry is a technique capable of high throughput analysis of cell suspensions with simultaneous measurement of physical and chemical properties of the cells [22], which has been extensively used for decades in clinical haematology to analyse blood cells. Flow cytometry constitutes three major components: fluidic circuitry for cell alignment, optical signal detection, and electronic conversion of digital signals into interpretable data. The acoustic-enhanced version of the flow cytometer instrument pre-focuses cells into a single stream using ultrasound waves before the interrogation point, resulting in much higher resolution than the hydrodynamic version [24]. Acoustic-enhanced flow cytometers can therefore be used successfully for small particle analysis such as bacteria.

We recently reported a flow cytometry assisted susceptibility testing (FAST) assay method to determine a carbapenem minimal inhibitory concentration in pure cultures of carbapenem-resistant *Klebsiella pneumoniae* using the nucleic acid intercalating dye, SYTO^®^ 9 [25]. This assay proved applicable to accurate detection of bacteria with the non-specific dye SYTO^®^ 9 in both unexposed and antibiotic-exposed bacteria in pure cultures. However, we believed the same approach could not be reliably used to detect bacteria in BC due to interference from blood-derived elements. Accordingly, we embarked on development of a novel diagnostic approach employing *in situ* hybridization with the bacteria-specific probe prior to acoustic flow cytometry. Here we describe development of the PNA-FISH enhanced acoustic flow cytometry assay that combines the advantages of PNA-FISH probes for fine molecular analysis with high throughput high-resolution cellular analysis of acoustic flow cytometry in order to increase bacterial detection specificity in blood culture and thus reduce the time to bacterial detection.

## Materials and methods

### Bacterial strains and culture media

Reference strains of *Klebsiella pneumoniae* (ATCC BAA 1705), *Pseudomonas aeruginosa* (ATCC 27853) and *Escherichia coli* (ATCC 25922) were used in the experiments described here. Bacterial stocks were prepared from bacteria by initial culture on blood agar (BA), then inoculation into Brain and Heart Infusion broth (BHIB) containing 10% glycerol and stored in −80°C freezer. These and other culture media including trypticase soy broth (TSB), Hanks Balanced Salt Solution (HBSS) and plate count agars (PCA) were obtained from the PathWest Laboratory Medicine WA (QEII, Nedlands, Western Australia). BacTEC aerobic blood culture bottles were obtained from Becton Dickinson (Franklin Lakes, New Jersey, USA). Fresh, uninfected human blood in EDTA was obtained from healthy adults on the day of the experiment in the specialist sample collection clinic of PathWest Laboratory Medicine WA (QEII, Nedlands, Western Australia) in accordance with institutional standard practice.

#### Fluorescent dye and probe

##### <u>Direct staining with</u> SYTO^®^ 9

One μL of SYTO^®^ 9 dye (ThermoFisher Scientific, Eugene, Oregon, USA) at a final concentration of 5μM was directly added to 1mL samples followed by a 5-minute incubation at room temperature before samples were analysed on the acoustic flow cytometer.

### PNA-FISH probe design

The eubacterial 16S probe (5’-Alexa488-O-TATCTAATCCTGTTT −3’) was designed using BioEdit software and NCBI BLAST (https://blast.ncbi.nlm.nih.gov/Blast.cgi). The peptide nucleic acid probe was conjugated to AlexaFluor 488 (supplied by PNA Bio Company, 107 N. Reino Rd., #242 Thousand Oaks, California, USA), and reconstituted prior to use in molecular grade water to 200μM stock concentration.

### Optimization of hybridization for pure bacterial cultures

The hybridization buffer was made in-house and contained 10% (w/v) dextran sulphate, 10mM NaCl, 0.1% (w/v) sodium pyrophosphate, 0.2% (w/v) polyvinylpyrrolidone, 0.2% (w/v) Ficoll, 5mM Na_2_EDTA, 0.1% (v/v) Triton X-100, 50mM Tris-HCl (pH 7.5). Wash buffer was also made in-house and contained 25mM Tris-HCl (pH 10.0), 137mM NaCl, and 3mM KCl. All reagents were obtained from Sigma Aldrich (St. Louis, Missouri, USA).

Three different concentrations of the PNA-FISH probe at 100 nM, 200 nM and 300 nM were used during optimization of hybridization. Other reaction parameters that were also optimized included the formamide (Sigma Aldrich, St Louis, Missouri, USA) concentration that varied from 0 - 50% (v/v), hybridization temperature that varied from 30°C - 60°C and duration of hybridization, which was either 15 or 30 minutes.

Pure bacterial cultures were grown overnight in 10mL TSB at 37°C without shaking and used for hybridization optimization where a 20μL aliquot was added to 480μL of hybridization buffer, followed by vortexing and incubation at varying temperatures and durations as described in figure legends. The sample was centrifuged at 10,000 *g* for 5 minutes and supernatant removed. The pellet was resuspended in a 500 μL wash solution and further incubated for 10 minutes at the same hybridization temperatures. The sample was centrifuged and washed once more in 500 μL wash solution, as described above. Samples were cooled to room temperature before they were analyzed on an acoustic flow cytometer (Attune NxT, ThermoFisher Scientific, Eugene, Oregon, USA). The PNA-FISH probe was added to give a final concentration of 200 nM in hybridization buffer containing formamide 30% (v/v), and was used in all subsequent procedures with simulated BC.

### Preparation of simulated BC

Simulated BCs were inoculated with bacteria grown in pure culture as follows: on day 1, stock bacteria were streaked onto the BA plates and incubated overnight at 37°C. On day 2, a single colony was picked, inoculated into pre-warmed TSB and incubated overnight at 37oC for 18 hours. On day 3, 1mL of overnight bacterial culture in TSB was pelleted by centrifugation at 10,000 *g* for 5 minutes followed by a wash and final resuspension in 1mL HBSS. An aliquot of bacterial suspension in HBSS was taken, diluted 1000-fold in HBSS and directly stained with 1 μL SYTO^®^ 9 for 5 minutes. The aliquot was analyzed on the acoustic flow cytometer (Attune, NxT, Thermo Fisher Scientific) to determine bacterial concentration.

Two aerobic BC bottles were prepared in parallel, each with 10 mL of uninfected human EDTA blood obtained from healthy adults. Subsequently, bacteria that were prepared and counted as described above were inoculated into the BC bottles at a final concentration of 10 CFU/mL. Following the BC bottle setup, one aliquot was taken immediately for plate count from one bottle. Both bottles were then incubated in the BacTEC BC autoanalyser. Aliquots were taken at 2-hour intervals for plate counts, PNA-FISH-AFC and other analyses from the same bottle sampled previously while the other bottle stayed untouched and was used for measuring a time to positivity.

### Hybridization protocol for simulated BC

Prior to hybridization, a lysis step was introduced for simulated BC to lyse blood cells and concentrate bacteria. Briefly, 1 mL of inoculated BC was harvested and added to 1 mL 10% Triton X-100 for 5 minutes followed by centrifugation at 10,000 *g* for 5 minutes. The pellet was re-suspended in 1 mL HBSS and centrifuged again followed by re-suspension of the pellet in 20 μL in HBSS. The sample was mixed with 480 μL of hybridization buffer, followed by vortexing and incubation at 40°C for 15 minutes. The sample was centrifuged at 10,000 *g* for 5 minutes and supernatant removed. The pellet was re-suspended in 500 μL wash solution and further incubated for 10 minutes at 40°C. The sample was centrifuged and washed once more in 500μL wash solution as described above. Samples were cooled to room temperature before they were analyzed by AFC.

Uninfected BC were lysed as described in the hybridization protocol for simulated BC above and stained with anti-CD 45 PerCP (ThermoFisher Scientific, Eugene, Oregon, USA) to identify blood elements of leukocyte origin by AFC analysis.

### Instrumentation and software

The acoustic flow cytometer (AFC) was used to analyze the samples. Flow cytometry analysis parameters are shown in Table 1. BacTEC FX blood culture system located in the Department of Microbiology at PathWest Laboratory Medicine WA was used to incubate inoculated blood cultures that rely on CO_2_ production as a measurement of bacterial growth. FlowJo software (FLOWJO, LLC, Oregon, USA) was used to analyze the data taken from the AFC. Spectral analysis of bacterial proteins was performed by Bruker MALDI Biotyper (Bruker Daltonics GmbH, Germany). PCR-based detection of bacteria in BC was performed on a FilmArray^®^ instrument (BioFire, version 1, BioMerieux Australia Pty Ltd, Baulkham Hills, NSW).

**Table 1.**
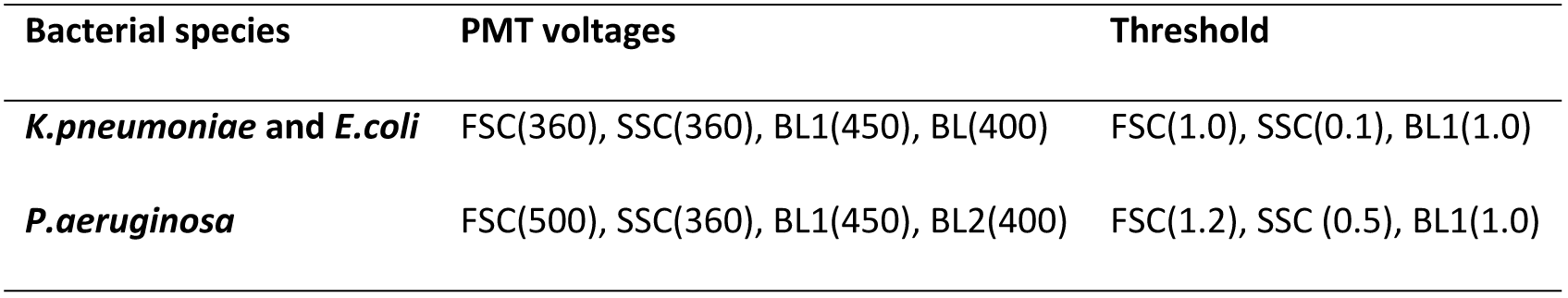
NxT AFC parameters.

| Bacterial species | PMT voltages | Threshold |
| --- | --- | --- |
| *K.pneumoniae* and *E.coli* | FSC(360), SSC(360), BL1(450), BL(400) | FSC(1.0), SSC(0.1), BL1(1.0) |
| *P.aeruginosa* | FSC(500), SSC(360), BL1(450), BL2(400) | FSC(1.2), SSC (0.5), BL1(1.0) |

### Event gating

The gating method for opto-electronic events produced by fluorescent dyes SYTO^®^ 9, and AlexaFluor 488 incorporated into a bacteria-specific PNA-FISH probe, was similar in pure bacterial cultures. Initially, the population of interest (POI) in the area of events corresponding to the bacterial population was determined by forward and side scatter (FSC-H/SSC-H) to allow detection of clusters of cells according to their size and density (Table 2). The doublet events, if present, were identified and excluded by the forward scatter (FSC-H/FSC-A). Single cells that stained positive with SYTO^®^9 or AlexaFluor 488 were further identified by their fluorescence intensity on the histogram. This population was then back-gated onto the FSC-H/SSC-H scatter plot to identify intact bacterial cells. This cell population represented pure single bacterial cells that incorporated the label (SYTO^®^ 9 or AlexaFluor 488) and was named ‘derived population of interest’ (dPOI). On all plots, dPOI was used for the final cell count.

**Table 2.**
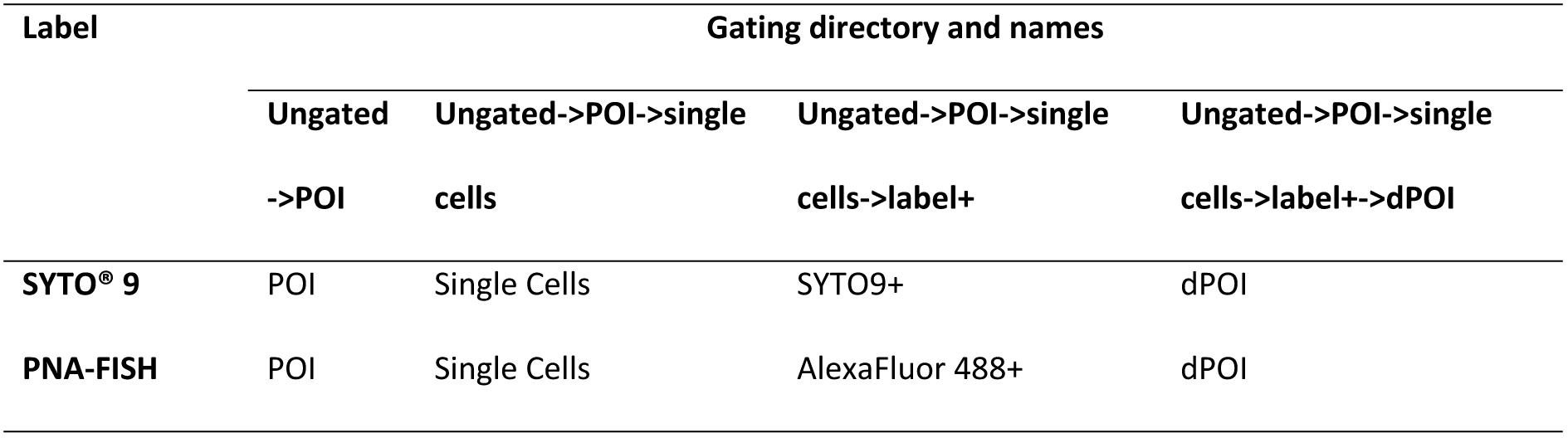
Gating methods flow chart.

## Results

### Bacterial detection by direct staining on AFC

Before developing a FISH-AFC-based assay for detection of bacterial DNA within bacterial cells using a specific probe, we first performed direct staining of pure bacterial cultures. To do this we used a non-specific nucleic acid dye, SYTO^®^ 9, and established instrument parameters as shown in Table 1. The gating method was developed as outlined in Material and Methods for bacterial cells directly stained with SYTO^®^ 9 (Fig 2A). Initially, a combination of forward scatter (FSC) versus side scatter (SSC) analysis was used to identify bacterial cells (so-called, population of interest, POI) based on their physical properties such as size and density (Fig 2A, i). This was followed by application of the FSC-H/FSC-A plot to distinguish intact single cells from doublets (Fig 2A, ii). Eventually, a subset of SYTO^®^ 9 positive (SYTO^®^ 9+) single cells was identified in a histogram (Fig 2A, iii) and used to pinpoint intact bacterial cells on the scatter plot by back-gating. This permitted ultimate identification of intact bacterial cells defined by their size and staining properties with SYTO^®^ 9 that were c alled a derived POI (dPOI) (Fig 2A, iv). When using direct staining with fluorescent dye SYTO^®^ 9 the limit of detection was determined as between 10^3^-10^4^ CFU/mL in pure culture (See Fig S1).

**Fig 2.**
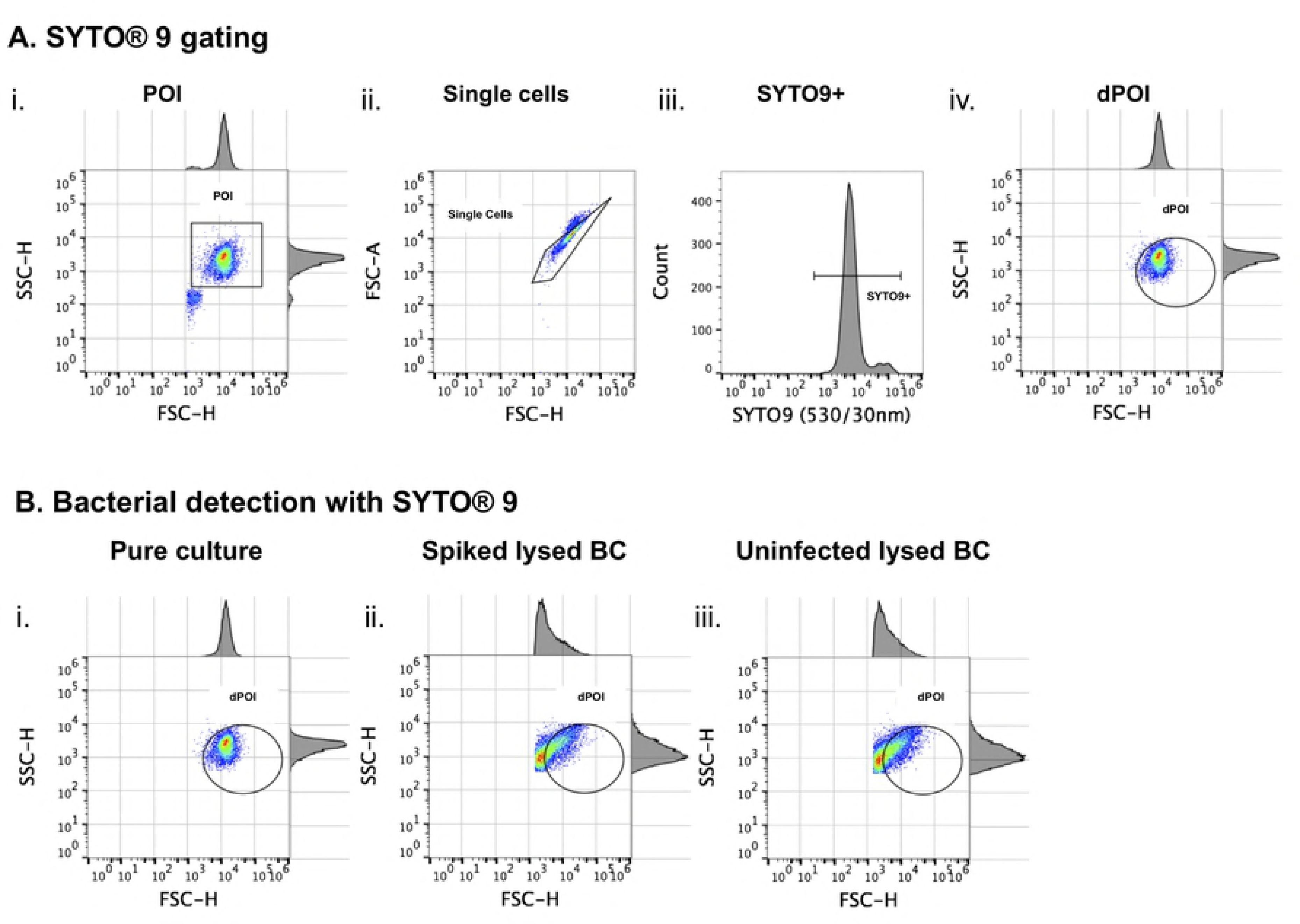
Bacterial detection by acoustic flow cytometry after direct staining with SYTO^®^ 9 in pure bacterial culture and inoculated blood culture. **A:** SYTO^®^ 9 gating for bacteria grown in TSB as pure culture: the population of interest (POI) was gated based on the FSC-H/SSC-H scatter plot (i) while single bacterial cell events were identified in the FSC-H/FSC-A plot (ii). Events that were SYTO^®^ 9 positive were gated in the BL1-H channel (iii) and back-gated to FSC-H/SSC-H scatter plot to identify bacterial cells labelled with SYTO^®^ 9, denoted as derived POI (dPOI) (iv) since SYTO^®^ 9 binds to other particles containing nucleic acids. **B:** Bacterial detection in pure culture (i) and blood culture (ii) by SYTO^®^ 9 staining. Uninfected BC was used as a negative control (iii). Both pure culture (i) and blood culture (ii) were inoculated with identical concentrations of bacteria at approximately 8.00 x 10^4^ CFU/mL. The pure bacterial culture was stained directly with SYTO^®^ 9, while both spiked and uninfected blood cultures were lysed with 10% Triton X-100 before SYTO^®^ 9 staining. Note the strong background noise produced during the BC lysis in both infected and uninfected BC samples overshadowing dPOI (ii and iii).

The next step was to test whether direct staining with SYTO^®^ 9 could be applied to detect bacteria in simulated blood cultures (BC). The simulated BC was spiked with similar bacterial concentration as in overnight pure culture at the time of detection, although due to its high viscosity and density it had to be either diluted or subjected to a lysis step before analysis on flow cytometer. Use of a dilution step for the BC was abandoned in favour of the lysis method due to reduced loss of bacterial cells and detection sensitivity. However, the lysis method produced an increase in background noise in the sample (Fig 2B). As shown in Fig 2B, the scatter plots of infected (Fig 2B ii) and uninfected (Fig 2B iii) BC following the lysis with 10% Triton X-100 were identical due to a non-specific binding of SYTO^®^ 9 dye to both bacterial cells and to abundant blood cell debris produced by lysis. Hence, detection of bacteria at low concentrations in BC could not be achieved by the direct staining method with SYTO^®^ 9.

### Development of *in situ* hybridization protocol

Due to the inability of SYTO^®^ 9 staining to discriminate bacteria from the background noise in BC samples, a unique bacteria-specific PNA-FISH probe hybridization protocol was developed. For this purpose, a eubacterial 16S PNA probe was designed with BioEdit software and labelled with the fluorophore AlexaFluor 488. Development of the fluorescence *in situ* hybridization (FISH) protocol required optimization of hybridization conditions such as formamide concentration, temperature, probe concentration and hybridization duration in order to obtain good signal-to-noise ratio and preserved cellular integrity. Accordingly, four optimization experiments were performed using pure bacterial cultures of *K.pneumoniae* as a model organism (Fig 3; Fig S1). Optimizations of formamide concentration and hybrization temperature were performed with SYTO^®^ 9 using the gating method that included both single cells and doublets in order to assess a recovery of the overall bacterial population stained with SYTO^®^ 9 (Fig S1). As shown in Fig 3, top two rows, 30% formamide in hybridization buffer and 40°C hybridization temperature provided optimal conditions for the best recovery of dPOI (Fig S2) in comparison to lower and higher formamide concentrations and hybridization temperatures, respectively.

**Fig 3.**
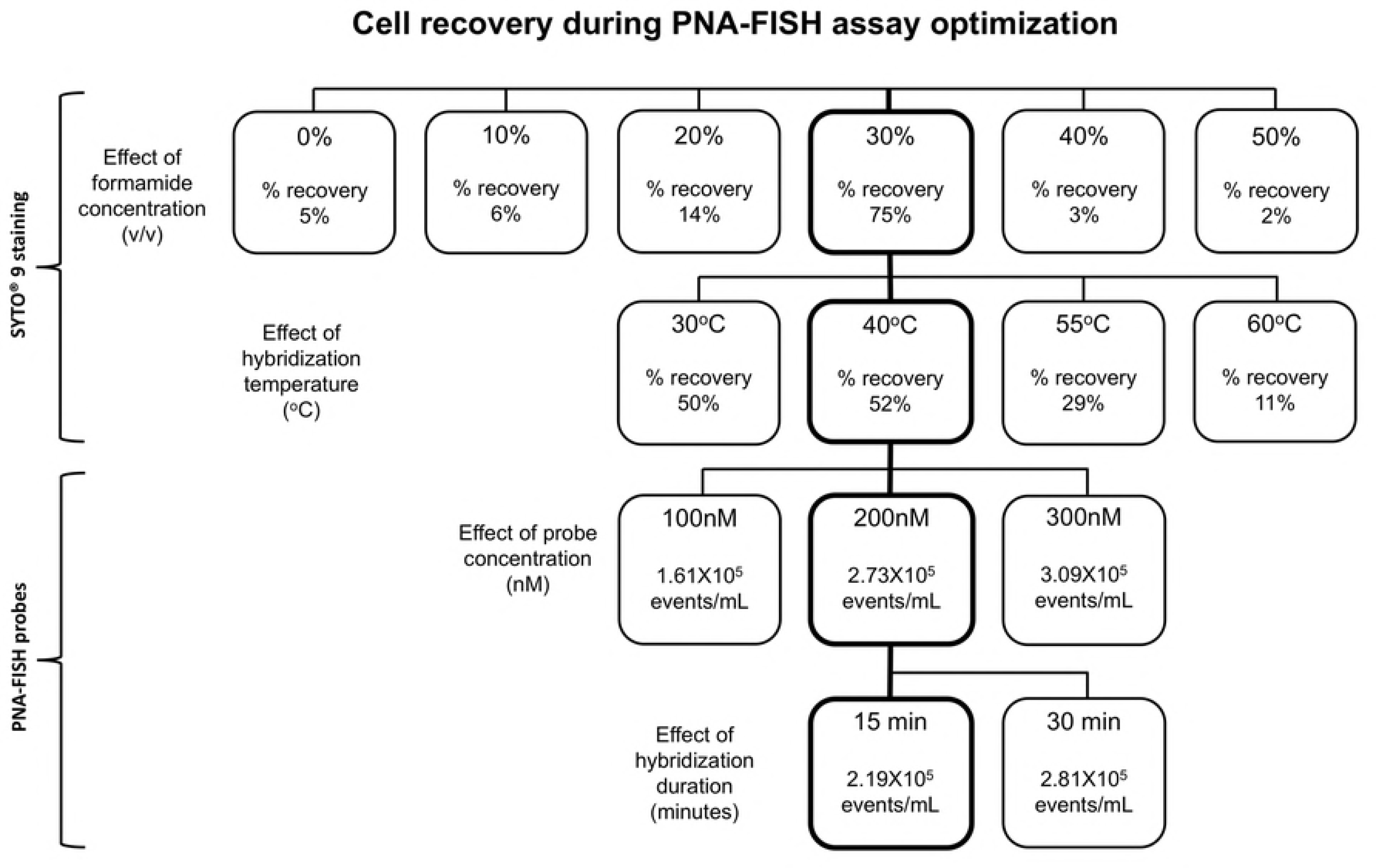
PNA-FISH assay optimisation flowchart using pure bacterial culture. Four optimisation protocols were performed one of each for formamide concentration, hybridization temperature, probe concentration, and hybridization duration. In the first two protocols (top rows), SYTO9 was used to determine changes in the overall bacterial population subjected to variations in formamide concentrations and temperature. Initially, varying concentrations of formamide were added to bacterial suspensions containing equal amounts of bacteria followed by incubation at 40°C for 15 minutes prior to staining with SYTO9. The highest bacterial recovery of 75% was achieved with 30% formamide (v/v, thick frame) compared to other treated and untreated samples. This formamide concentration (30%) was selected for subsequent experiment in which bacterial cells were exposed to varied hybridization temperatures from 30°C to 60°C (second row). The best result with 52% bacterial cell recovery was obtained at 40°C (thick frame). For the last two optimisation steps, PNA-FISH probe was used in hybridization buffer containing 30% formamide at 40°C (third and fourth rows). In the third row, three different probe concentrations from 100 to 300 nM were tested by PNA-FISH. The probe concentration of 200 nM turned out to be optimal (thick frame) since no significant increase in hybridisation signal was observed when the probe concentration was elevated to 300 nM. Eventually, hybridization duration of 15 minutes was determined as optimal since the majority of the hybridization positive events were detected in the first 15 minutes. The squares with thick frames stacked vertically indicate optimised hybridization conditions applied to all subsequent experiments.

The hybridization signal was optimised using PNA-FISH probe as described in Fig S1B, altering two assay parameters, PNA-FISH probe concentration from 100nM to 300nM, and hybridization duration, from 15 to 30 minutes (Fig 3). As shown in Fig 3 and Fig S2, optimal hybridization signal intesity was obtained with probe concentration at 200nM following 15 minutes of hybridization. In summary, optimal conditions for PNA-FISH hybridization were determined to include 30% formamide (v/v) and 200 nM probe in hybridization buffer, 40°C hybridization temperature and 15 minutes incubation time (Fig 3; flowchart squares with thick frames). These conditions were applied to all subsequent PNA-FISH experiments.

### PNA-FISH gating method for BC

A gating method for the PNA-FISH assay in BC differed from the gating method applied previously to PNA-FISH in pure bacterial culture (Fig S1B) due to an overlap between blood and probe specific signals in the 530/30nm channel (data not shown). A different approach was developed for the PNA-FISH in BC based on the difference between the specific signal of the AlexaFluor 488-labeled PNA-FISH probe attached to the bacterial DNA (530/30nm) and a non-specific autofluorescence (574/26nm) of the lysed blood. Accordingly, the bacterial population could be found in the BL1 channel (AlexaFluor 488) while the lysed blood elements (autofluorescence) were observed in the BL2 channel (Fig 4, i). The bacterial population was further refined by the FSC-H/FSC-A scatter plot that was used to gate out the doublets (Fig 4, ii). Distribution of signal intensity of the AlexaFluor 488 on the histogram indicated uniform probe hybridization (Fig 4, iii). This was later used as a bacteria-specific signal to gate events representing pure bacterial cell population on the FSC-H/SSC-H scatter plot that was labeled as dPOI (Fig 4, iv).

**Fig 4.**
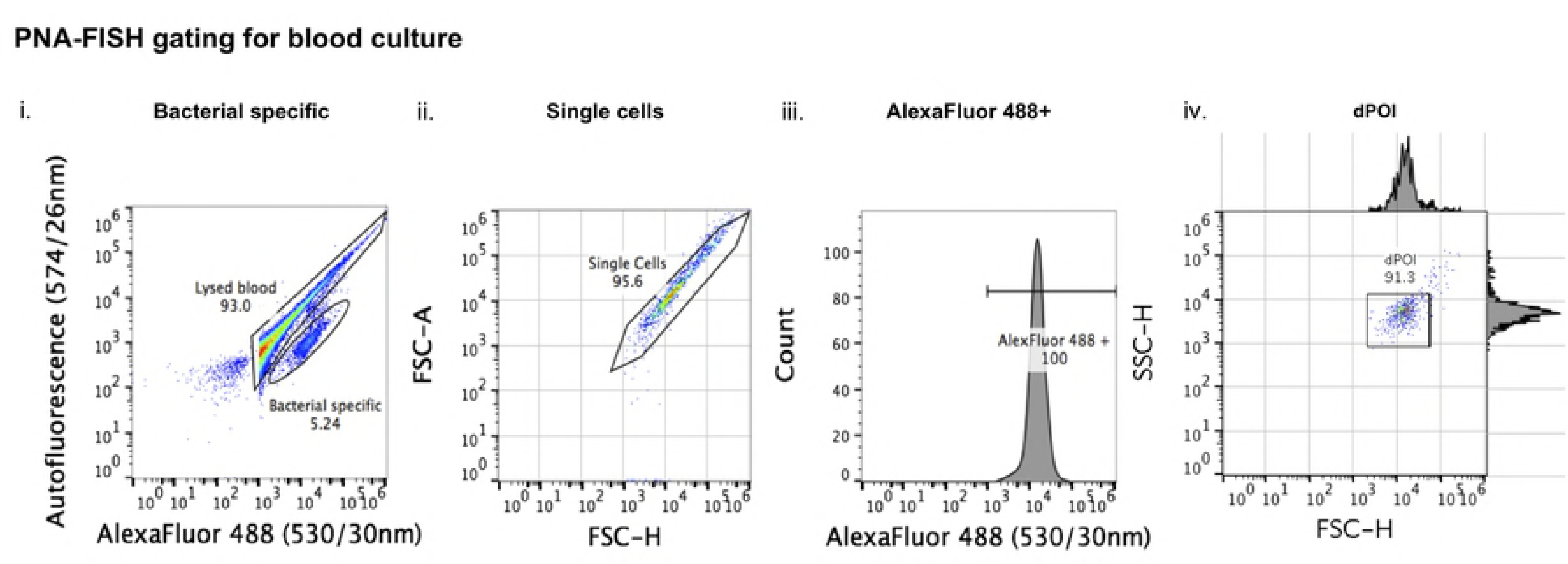
Gating of PNA-FISH hybridized BC samples. Bacteria-specific events were separated from the background noise of lysed blood based on a difference between the PNA probe-specific signal and autofluorescence produced by blood (i). This was followed by identification of single cells on the FSC-H/FSC-A scatter plot within the bacterial population (ii). Probe specific signal was gated as AlexaFluor 488 + on the histogram (iii), and all events within this gate were backgated and shown on the FSC-H/SSC-H plot as dPOI (iv).

The validity of gating off blood debris using autofluorescence was confirmed in a separate experiment in which the white blood cell-specific antibody anti-CD45 PerCP was used. This antibody bound to the same population observed by autofluorescence but not to pure bacterial population confirming that autofluorescence originated from the lysed blood (data not shown).

### Bacterial detection in BC by PNA-FISH enhanced AFC

A pair of aerobic bottles spiked with approximately 10 CFU/mL of bacteria were set up for three different bacterial species, *E. coli, K. pneumoniae and P. aeruginosa* and incubated in the BacTEC system. Aliquots (1mL) from one bottle of the pair were removed hourly, treated with 10% Triton X-100 and the bacteria concentrated and washed by centrifugation. The samples were then hybridized with the PNA-FISH probe under optimized conditions and analyzed on the NxT AFC. The other BC bottle remained incubating on the BacTEC system until the positivity threshold had been reached.

Using *E. coli* as a test species for PNA-FISH flow cytometry, the first positive signal was detected at 5 hours of incubation following hourly sampling and PNA-FISH analysis within the bacteria-specific gate (Fig 5A). The number of events within the bacteria-specific gate increased further after 6 and 7 hours of incubation, indicating actively growing bacteria (Fig 5A). This experiment was repeated three times and time to detection (TTD) with PNA-FISH-AFC for *E. coli* in the spiked BC was determined to be between 5-6 hours. Simultaneously, growth of *E. coli* in spiked BC was monitored by plate count at 2-hour intervals from the same BC bottle (Fig 5B). The time to blood culture-based positivity for *E. coli* was determined in the separate, simultaneously incubated BC bottle to be between 13-14 hours with bacterial concentrations reaching values from 4.59 x10^8^ to 9.20 x 10^8^ CFU/mL at this point (Fig 5B).

**Fig 5.**
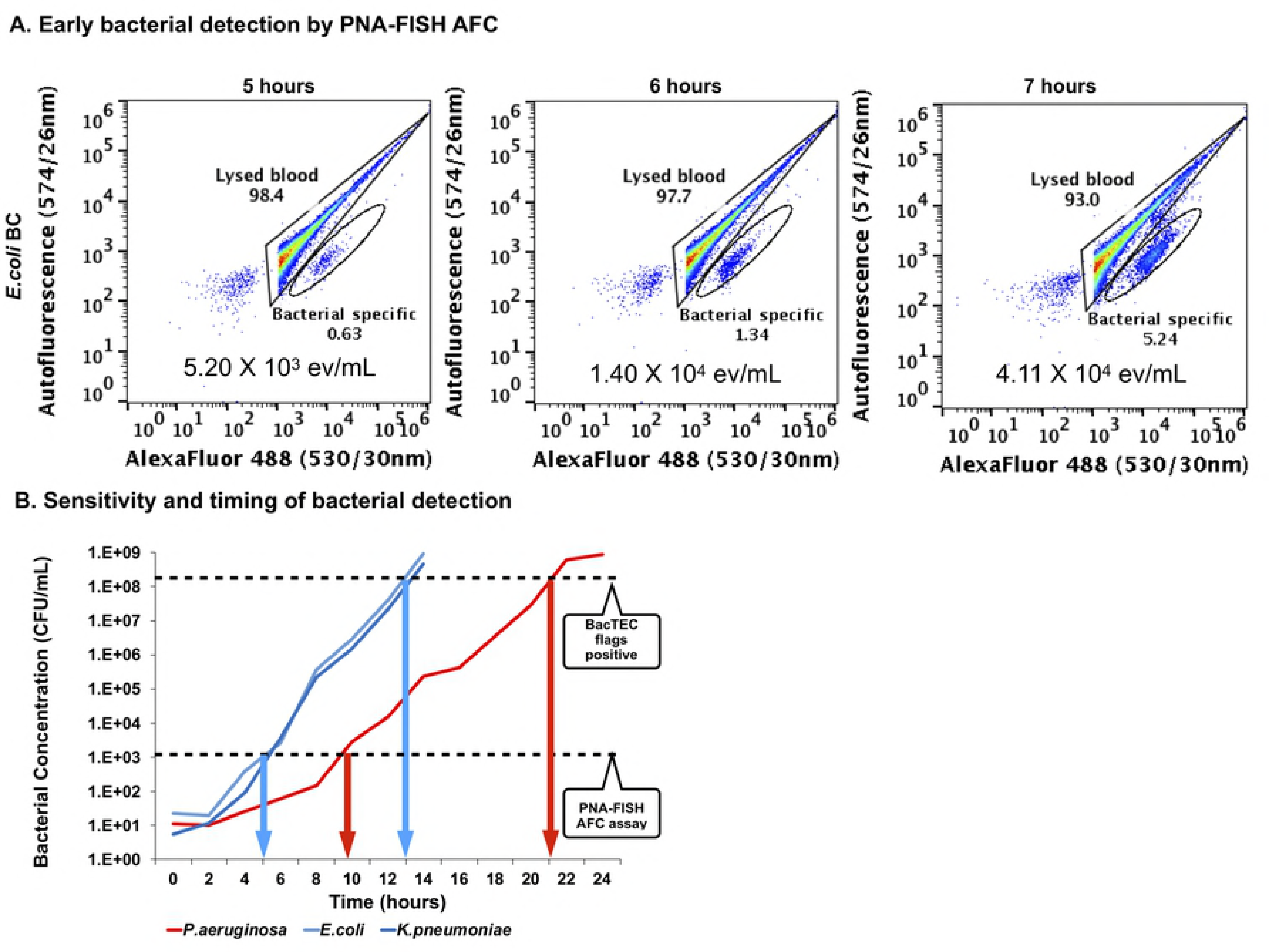
Early bacterial detection in BC with PNA-FISH-AFC. **A:** Detection of *E. coli* by PNA-FISH enhanced AFC in spiked BC following incubation on the BacTEC FX system. A 9 time increase in number of events was observed in the bacteria-specific gate from 5 to 7 hours of incubation indicating actively growing bacterial population. **B:** Growth curves of bacteria in spiked BC on the BacTEC system. Two fast growing bacterial species, *E coli* and *K pneumoniae*, and one slow grower, *P aeruginosa*, were incubated on the BacTEC FX system. The 2-hourly determined plate counts (*y*-axis, CFU/mL) were used to produce growth curves, while the appearance of bacteria-specific events by PNA-FISH enhanced AFC was monitored hourly for the first 12 hours of incubation. The overall positivity on the BacTEC system was also monitored in parallel. Bacteria-specific signal by PNA-FISH-AFC was first observed at 5 and 10 hours of incubation for the fast (*E. coli, K. pneumoniae*) and slower growing bacterial species (*P. aeruginosa*), respectively. This was confirmed by three independent experiments. The positivity on the BacTEC system was reached at approximately 13 hours and 21 hours of incubation for the fast and slow growers, respectively, also confirmed by three independent experiments with slight variations between experiments. The horizontal broken lines indicate the sensitivity of the PNA-FISH-AFC at 10^3^-10^4^ CFU/mL, and for the BacTEC system at 108-109 CFU/mL. The time gained by the PNA-FISH assay when compared to BacTEC positivity was 8 hours for fast growers, as indicated by blue arrows, and 10 hours for slow grower, as indicated by red arrows, respectively.

Similar TTD results were obtained for *K. pneumoniae* using PNA-FISH-AFC analysis (see Fig S3). However, the first positive signal for *P. aeruginosa* by PNA-FISH-AFC was observed after 10 hours of incubation in the BacTEC system (Fig S3). Combined analyses for all three-bacterial species revealed that the first positive signal by PNA-FISH-AFC occurred at a bacterial concentration of 10^3^-10^4^ CFU/mL reached by *E. coli* and *K. pneumoniae* after 5 hours in BC and after 10 hours in BC by *P. aeruginosa*, respectively (Fig 5B). The time to positivity of *E. coli* and *K. pneumoniae* were similar on the BacTEC in the range of 13-14 hours of incubation, while the growth of *P. aeruginosa* required 19-21 hours to reach the arbitrary levels due to a slower growth rate (Fig 5B). Since slow growers showed a less steep gradient than fast growers, there was more time gained to detect slow-growing organisms in blood culture when the PNA-FISH assay was applied (Fig 5B).

### Bacterial enumeration - PNA-FISH-AFC *versus* plate count

Each aliquot of spiked BC that was analyzed by PNA-FISH flow cytometry was also enumerated by the plate count method for comparison as indicated in Fig 5B. Biological triplicates were set up for both assays and average bacterial concentrations determined by either PNA-FISH or plate counts are presented in Table 3. As shown on Table 3, the number of events determined by PNA-FISH-AFC was lower than the number of colony forming units (CFU) determined by plate count. Plausible explanations include cell loss during hybridization due to multiple washes and exposure to high temperatures, as well as application of gating methods that removed the bacterial doublets during post analysis. Probe efficiency was very high (97.1%) as expected of peptide nucleic acid probes, hence this did not contribute to the cell loss.

**Table 3.** Bacterial enumeration using PNA-FISH and plate counts at the time of detection.

| Bacterial species | BC incubation in the BacTec system (hours) | PNA-FISH-AFC (events/mL±SD) | Plate counts (CFU/mL ± SD) |
| --- | --- | --- | --- |
| *Escherichia coli* | 5 | $0.17 \times 10^4 \pm 0.11$ | $0.21 \times 10^4 \pm 0.27$ |
| | 6 | $0.40 \times 10^4 \pm 0.26$ | $0.57 \times 10^4 \pm 0.72$ |
| | 7 | $0.28 \times 10^4 \pm 2.72$ | $2.94 \times 10^4 \pm 1.05$ |
| *Klebsiella pneumoniae* | 5 | $0.25 \times 10^4 \pm 0.40$ | $0.44 \times 10^4 \pm 0.62$ |
| | 6 | $0.64 \times 10^4 \pm 0.79$ | $1.41 \times 10^4 \pm 1.44$ |
| | 7 | $5.29 \times 10^4 \pm 6.24$ | $9.24 \times 10^4 \pm 12.2$ |
| *Pseudomonas aeruginosa* | 10 | $1.09 \times 10^4 \pm 0.80$ | $1.31 \times 10^4 \pm 1.38$ |
| | 11 | $2.36 \times 10^4 \pm 1.81$ | $3.10 \times 10^4 \pm 3.80$ |
| | 12 | $7.52 \times 10^4 \pm 8.31$ | $12.1 \times 10^4 \pm 18.2$ |

### Performance of PNA-FISH-AFC relative to MALDI-TOF and FilmArray^®^

Two additional culture-independent assays were set up in parallel with PNA-FISH-AFC, MALDI-TOF and FilmArray^®^, that are routinely used in clinical laboratories. When all three assays were performed in parallel, the TTD using PNA-FISH-AFC was shorter than using MALDI-TOF for all three organisms; FilmArray^®^ and PNA-FISH-AFC were equally sensitive for *K. pneumoniae* and *P. aeruginosa* resulting in similar TTD; while PNA-FISH-AFC was more sensitive than FilmArray^®^ for *E. coli* shown by 2-hours shorter TTD (Table 4). This experiment was performed three times, demonstrating consistently shorter TTD by PNA-FISH-AFC than MALDI-TOF for all three bacterial species and a limited advantage of PNA-FISH-AFC over FilmArray^®^ assays for *E coli*.

**Table 4.**
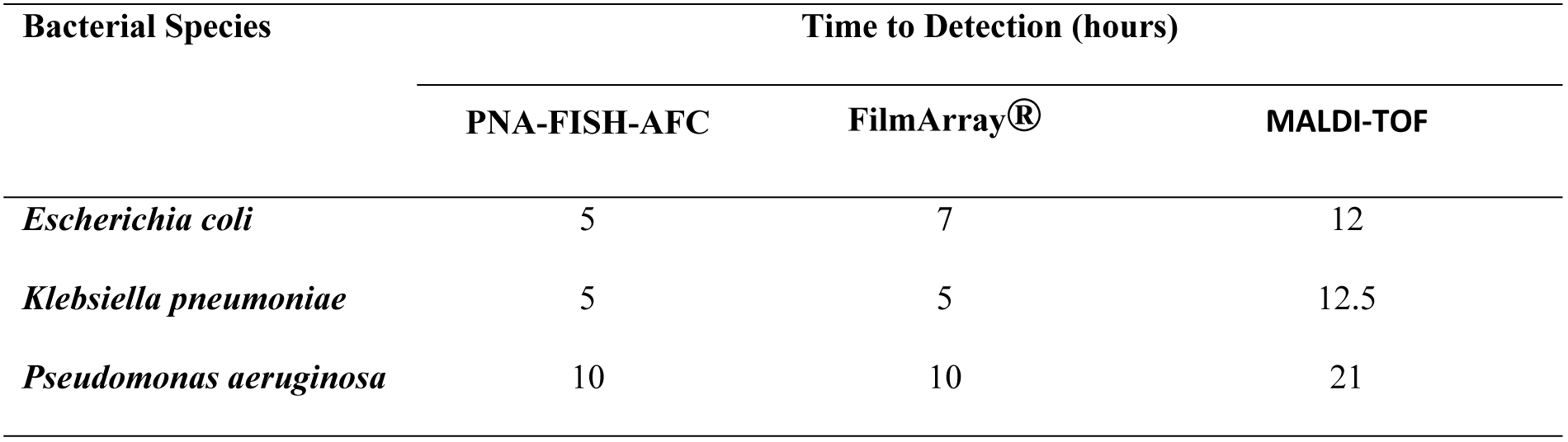
Earliest time to detection using different diagnostic assays.

| Bacterial Species | Time to Detection (hours) |  |  |
| --- | --- | --- | --- |
|  | PNA-FISH-AFC | FilmArray® | MALDI-TOF |
| *Escherichia coli* | 5 | 7 | 12 |
| *Klebsiella pneumoniae* | 5 | 5 | 12.5 |
| *Pseudomonas aeruginosa* | 10 | 10 | 21 |

## Discussion

Since the early 20^th^ century there has been a search for faster ways to detect bacteria in the bloodstream. While direct detection has thus far been unachievable due to low bacterial starting concentrations of 1-10 CFU/mL of blood, a universal blood culture system that could support growth and increase the concentration of any clinically significant bacterial species has been developed. Unfortunately, nutritional requirements for bacterial species differ, so does the time to achieve a minimum concentration that would give reliable results in subsequent diagnostic tests, ranging from several hours to several days in culture. Bacterial concentrations in BCs at the time of positivity in autoanalyser systems have been established at 10^7^-10^9^ CFU/mL [13-23]. These figures are in agreement with our data from the BacTec system at the time of positivity, and determined by both flow cytometry and plate counts to be in the range of 4.59-9.20X10^8^ CFU/mL for three commonly isolated bacterial species, *E. coli, K. pneumoniae* and *P. aeruginosa*.

The introduction of MALDI-TOF analysis has significantly improved the speed of bacterial identification from positive BCs. However, MALDI-TOF relies on a high bacterial concentration and still requires extended incubation time in BCs. It may also require a purity subculture, particularly in polymicrobial and streptococcal bacteraemias where the subculture is mandatory [26, 27]. Recently, a pre-lysis step was introduced prior to bacterial identification by MALDI-TOF in order to avoid the need for subculture [28-31]. This has reduced the turnaround time to bacterial identification to approximately 2 hours after the BC bottle flagged positive although the BC time to positivity had to be extended to allow sufficient bacterial concentration for MALDI-TOF analysis [22].

Here we describe a novel approach to bacterial detection with peptide nucleic acid (PNA) fluorescence *in situ* hybridization (FISH) enhanced acoustic flow cytometry (eAFC) that is applied directly to positive BCs at earlier time point without the subculture step. The use of a eubacterial 16S rRNA probe specifically targets bacterial nucleic acids while the flow cytometer determines cellular characteristics of bacteria from their physical and fluorescence properties. In contrast to the PCR-based approaches that do not distinguish between genetic material that is either cell-free or within bacterial cells, the combination of molecular and cellular analyses in PNA-FISH-AFC specifically detects genetic material within intact bacterial cells. Thus, the integrity of bacterial cells becomes another diagnostic feature of this assay that indicates the presence of bacteria in a patient’s bloodstream.

The fluorophore selected for the FISH assay produces a signal within the green spectrum where much of the autofluorescent background noise produced by leukocytes was observed [32]. Indeed, leukocytic debris was labeled with anti-CD45 antibody conjugated to PerCP in lysed uninfected and spiked BC. In contrast, pure bacterial cultures were CD45 negative (data not shown). The gating method using 530/30nm and 574/26nm scatter plot on the NxT acoustic flow cytometer we developed here proved effective in discriminating lysed blood elements from the bacterial population. This has significantly increased the sensitivity of bacterial detection in BC to the threshold values of 10^3^ – 10^4^ CFU/mL and decreased time to positivity to only 5 hours for rapidly dividing *K. pneumoniae* and *E.coli*, and 10 hours for slow growers such as *P. aeruginosa*.

The newly developed PNA-FISH assay proved more sensitive than MALDI-TOF analysis for detecting three bacterial species in BC, namely *E. coli*, *K. pneumoniae* and *P. aeruginosa,* by four to five orders of magnitude. Here, we used a universal eubacterial probe to optimize the assay. In future, we envisage the use of bacterial species-specific probes for the most frequent bacterial species associated with sepsis.

The time to detection of FilmArray^®^ was similarly effective as PNA-FISH for early detection of *K. pneumoniae* and *P. aeruginosa,* however, this technology is PCR-based and provides evidence only for the presence of bacterial nucleic acids. In the case of *E. coli* detection, FilmArray^®^ in common with other PCR-based assays, shows decreased sensitivity due to a need to avoid false positives as it is a common contaminant of laboratory reagents. A key advantage of the latter is the detection of bacterial nucleic acids within intact bacterial cells despite a similarity in the time to detection between FilmArray^®^ and PNA-FISH-AFC, and the ability of both methods to provide molecular identification.

A further advantage of our assay method is increased specificity and sensitivity of detection of intact bacterial cells in BC that can reduce time to bacterial detection by 6 to 12 hours. The optimized protocol allows simultaneous set up of multiple samples with a simple procedure for manual sample acquisition and processing. Overall, it takes 1.5 hours to complete the FISH step with multiple samples, and an additional 3 minutes for AFC reading per sample. In future, we plan to use multiple species-specific probes for early bacterial identification in polymicrobial infections.

The novel diagnostic assay, PNA-FISH-AFC, described here has a potential to significantly reduce the turnaround time of bacterial detection and identification in blood culture in large clinical laboratories, and may allow blood culture autoanalyser systems to use lower threshold values for earlier detection of bacteraemia. In septicaemic patients with high bacterial loads in peripheral blood, this novel assay has the potential to identify the causative agent(s) directly from the whole blood. Ultimately, the major potential of PNA-FISH-AFC is in early and enhanced bacterial detection in blood culture that will improve survival and management of bacteraemic patients.

## Competing interests

XXH receives a PhD scholarship from PathWest Laboratory Medicine WA; TJJI received in-kind support from Thermo Fisher Scientific in the form of loan equipment (acoustic flow cytometer): http://corporate.thermofisher.com/en/home.html

## Ethics statement

The study was conducted in accordance with the Australian National Health Research Ethics guidelines (NH&MRC), the Declaration of Helsinki and Good Clinical Practice guidelines. Anonymised samples without identifiable clinical data can be used for *in vitro* pathology test development without further Health Research Ethics Committee approval. The work was conducted as a clinical laboratory quality initiative, in which no patient samples were used. The authors did not collect the blood, which was obtained from healthy adult volunteers by the state pathology service’s phlebotomists at a designated specimen collection centre. Institutional governance requirements were met by completing a specimen collection request form on each occasion volunteer blood samples were required.

## Acknowledgement

We would like to thank Mr Adam Merritt from the Molecular Diagnostics Laboratory at PathWest in QEII Medical Centre for his assistance with the design of the PNA-FISH probe and Dr Christine Carson for critical reading of the manuscript.

## Supplementary Data

**S1. Gating method for PNA-FISH optimization experiments on pure *K.pneumoniae* culture. A**: SYTO^®^ 9 was used in the formamide and hybridization temperature optimization experiments to investigate the overall bacterial population that contained both single and multiple events. POI was identified based on FSC-H/SSC-H plots, from which SYTO^®^ 9 positive population was identified on the histogram. A 10% contour was applied to the SYTO^®^ 9 positive population and labelled derived POI (dPOI). **B:** PNA-FISH gating on pure bacterial culture where POI was identified based on FSC-H/SSC-H plot followed by FSC-H/FSC-A to exclude doublet events. Probe specific events were identified from the singlet population as AlexaFluor 488+ on the histogram and this population was further gated as dPOI using FSC-H/SSC-H plot.

**S2. Flow charts of PNA-FISH-AFC optimization.** Four optimization experiments were set up sequentially to identify optimal conditions for bacterial detection by PNA-FISH-AFC as shown in Figure 3. Thick frames indicate the optimal conditions selected for subsequent experiments. Firstly, bacteria in pure culture were exposed to varying concentrations of formamide. The effect of increasing concentrations of formamide from 20 to 40% (v/v) on *K. pneumoniae* was monitored by SYTO^®^ 9 staining and presented as dPOI in the FSC-H/SSC-H plots (top row). The best cell preservation with tight bacterial population in the dPOI was observed in 30% formamide when compared to buffers containing 20 and 40% (v/v) formamide. In the second row, bacteria were hybridized in buffer containing 30% formamide (v/v) for 15 minutes at varied hybridization temperatures of 30°C, 40°C and 55°C followed by SYTO^®^ 9 staining. At temperatures of 30°C and 40°C, a better cell preservation was observed with minimal debris to the left of dPOI than at 55°C, where the debris subpopulation to the left of dPOI increased indicating likely degradation of bacterial cells at elevated temperature. In a third row, bacteria were hybridized at 40oC for 15 minutes in hybridization buffer containing 30% formamide (v/v) followed by two washes using wash buffer. Signal intensities of PNA-FISH probe concentrations of 200 nM and 300 nM were compared in a histogram where no significant difference between them was observed. Hence, 200 nM probe concentration was selected for the following experiment to test hybridization duration. In the last experiment (bottom row), two aliquots with the same concentration of bacteria were subjected to PNA-FISH assay for either 15 or 30 minutes under the conditions described above. The signal intensity was 1 log stronger after 15 minutes hybridization (red) than after 30 minutes (blue). Hence, the optimal hybridization condition for PNA-FISH-AFC was determined in the presence of 30% formamide (v/v) and 200 nM probe at 40oC for 15 minutes.

**S3. Early bacterial detection in spiked BC with *K. pneumoniae* and *P. aeruginosa.*** Similar to experiments with *E. coli* spiked BC (Fig 5), two other bacterial species, *K. pneumoniae* and *P. aeruginosa* were subjected to PNA-FISH-AFC following extended incubation in BC. The lowest bacterial concentration detected for both species was between 10^3^-10^5^ ev/mL as shown below the bacteria-specific population on the plots. While *K. pneumoniae* was first detected by PNA-FISH at 5 hours of incubation on the BacTEC FX system (top row), it took 10 hours to detect *P. aerugin*osa (bottom row). Additional 2 hours of incubation resulted in an increase of bacteria specific population for both organisms.

